# A Single-Cell Network Approach to Decode Metabolic Regulation in Gynecologic and Breast Cancers

**DOI:** 10.1101/2024.09.18.613640

**Authors:** Akansha Srivastava, P K Vinod

**Affiliations:** Centre for Computational Natural Sciences and Bioinformatics, IIIT, Hyderabad-500032, India

**Keywords:** single-cell transcriptomics, gynaecologic cancers, malignant heterogeneity, metabolic pathways, transcriptional regulation

## Abstract

Cancer metabolism is characterized by significant heterogeneity, presenting challenges for treatment efficacy and patient outcomes. Understanding this heterogeneity and its regulatory mechanisms at single-cell resolution is crucial for developing personalized therapeutic strategies. In this study, we employed a single-cell network approach to characterize malignant heterogeneity in gynecologic and breast cancers, focusing on the transcriptional regulatory mechanisms driving metabolic alterations. By leveraging single-cell RNA sequencing (scRNA-seq) data, we assessed the metabolic pathway activities and inferred cancer-specific protein-protein interactomes (PPI) and gene regulatory networks (GRNs). We explored the crosstalk between these networks to identify key alterations in metabolic regulation. Clustering cells by metabolic pathways revealed tumor heterogeneity across cancers, highlighting variations in oxidative phosphorylation, glycolysis, cholesterol, fatty acid, hormone, amino acid, and redox metabolism. Our analysis identified metabolic modules associated with these pathways, along with their key transcriptional regulators. Notably, transcription factors related to ER stress, immune response, and cell proliferation, along with hypoxia-inducible factor and sterol regulatory element-binding proteins were found to drive metabolic reprogramming. These findings provide new insights into the complex interplay between metabolic rewiring and transcriptional regulation in gynecologic and breast cancers, offering potential avenues for targeted therapeutic strategies in precision oncology. Furthermore, this pipeline for dissecting coregulatory metabolic networks can be broadly applied to decipher metabolic regulation in any disease at single-cell resolution.

## Introduction

Gynecological malignancies, primarily encompassing endometrial, ovarian, and cervical cancers, significantly impact women’s health on a global scale^1^. These cancers originate from the Müllerian ducts, sharing similar embryonic origins. Additionally, breast cancer is the most commonly diagnosed cancer among women and the leading cause of cancer deaths in women worldwide^2^. Breast cancer is categorized into four subtypes based on the expression of hormone receptors: estrogen receptor positive (ER+), progesterone receptor positive (PR+), human epidermal growth factor receptor positive (HER2+), and triple-negative breast cancer (TNBC). ER+ breast cancer expresses the estrogen receptor, while TNBC lacks expression of ER, PR, and HER2, making it unresponsive to hormonal therapies. Recent TCGA pan-cancer studies have revealed molecular similarities across gynecologic and breast cancers, highlighting both common pathways and unique molecular features^3,4^. These cancers pose significant health challenges due to their prevalence and the complexity of their diagnosis and treatment. Despite advancements in medical research, gynecological cancers are often diagnosed at advanced stages, resulting in poor prognoses and high mortality rates^1^. In particular, ovarian cancer is frequently diagnosed at a late stage, contributing to its high mortality rate and poor survival outcomes^5^.

Single-cell RNA sequencing (scRNA-seq) has revolutionized cancer research by allowing the examination of gene expression at the individual cell level. This technology provides insights into the heterogeneity of cancer cells within a tumor, revealing distinct cellular subpopulations and their functional states. Differential expression analysis of scRNA-seq data can reveal genes unique to specific cell types and states. However, deciphering cellular functions based solely on lists of upregulated or downregulated genes is challenging^6^. The functional roles of genes and the impact of disease-associated variants are significantly influenced by their interaction partners within the cellular context. Network modeling of gene interactions helps elucidate the functional organization of key regulators and the pathways they govern in each cell state^6^. This approach has shifted our perception of cellular mechanisms from linear pathways to a complex web of molecular interactions^6^. A recent study analyzed cell type-specific gene expression changes to identify potential drug targets and genetic biomarkers associated with Alzheimer’s disease using single network biology^7^. By constructing and analyzing networks of genes, we can elucidate the regulatory circuits that drive cancer progression.

Cancer metabolism has gained significant attention in the last decade due to its crucial role in supporting tumor growth and survival. Cancer cells often reprogram their metabolic pathways to meet the demands of rapid proliferation and adaptation to the tumor microenvironment. A hallmark of this metabolic reprogramming is the Warburg effect, where cancer cells preferentially utilize aerobic glycolysis over oxidative phosphorylation, even in the presence of adequate oxygen. This metabolic shift not only provides energy but also supplies the building blocks necessary for macromolecule biosynthesis. The transcriptional regulation of metabolism in cancer involves complex networks of transcription factors that control the expression of genes related to various metabolic pathways. Investigating the intricate regulation of metabolic pathways in cancer cells can uncover novel targets for cancer therapy and deepen our understanding of tumor biology.

Earlier studies on metabolism focused on analyzing the tissue-level changes. The field of single-cell transcriptomics provides the opportunity to study metabolism at the single-cell level and it is an emerging area of research. Few studies have utilized scRNA-seq to investigate metabolic changes during developmental processes^8^. Different computational modeling approaches such as pathway-level analysis, constraint-based modeling, and kinetic modeling can be adopted to study single-cell metabolism^9^. Pathway-level analysis focuses on identifying metabolic pathways and their activity levels within individual cells by integrating transcriptomic data with prior knowledge of metabolic networks. In constraint-based modeling, cellular metabolism is represented as a network of reactions with constraints based on known biochemical principles to predict metabolic fluxes and identify active pathways. Kinetic models describe the dynamics of individual reactions and enzyme kinetics. A recent study adopted a pathway-based approach to establish a computational framework for characterizing metabolism using scRNA-seq data, shedding light on the unique metabolic demands of different cell types within the tumor microenvironment^10^.

In this work, we employed a scRNA-seq network biology approach to investigate the metabolic landscape and its regulation in gynecologic and breast cancers. Figure 1 illustrates the workflow of our study. We examined the crosstalk between different cellular networks to uncover the coregulatory networks involved in metabolic reprogramming. We have created a comprehensive resource on single-cell cancer metabolism. This resource offers valuable insights into regulatory mechanisms at play and contributes to the advancement of personalized medicine for treating gynecologic and breast cancers.

**Figure 1:**
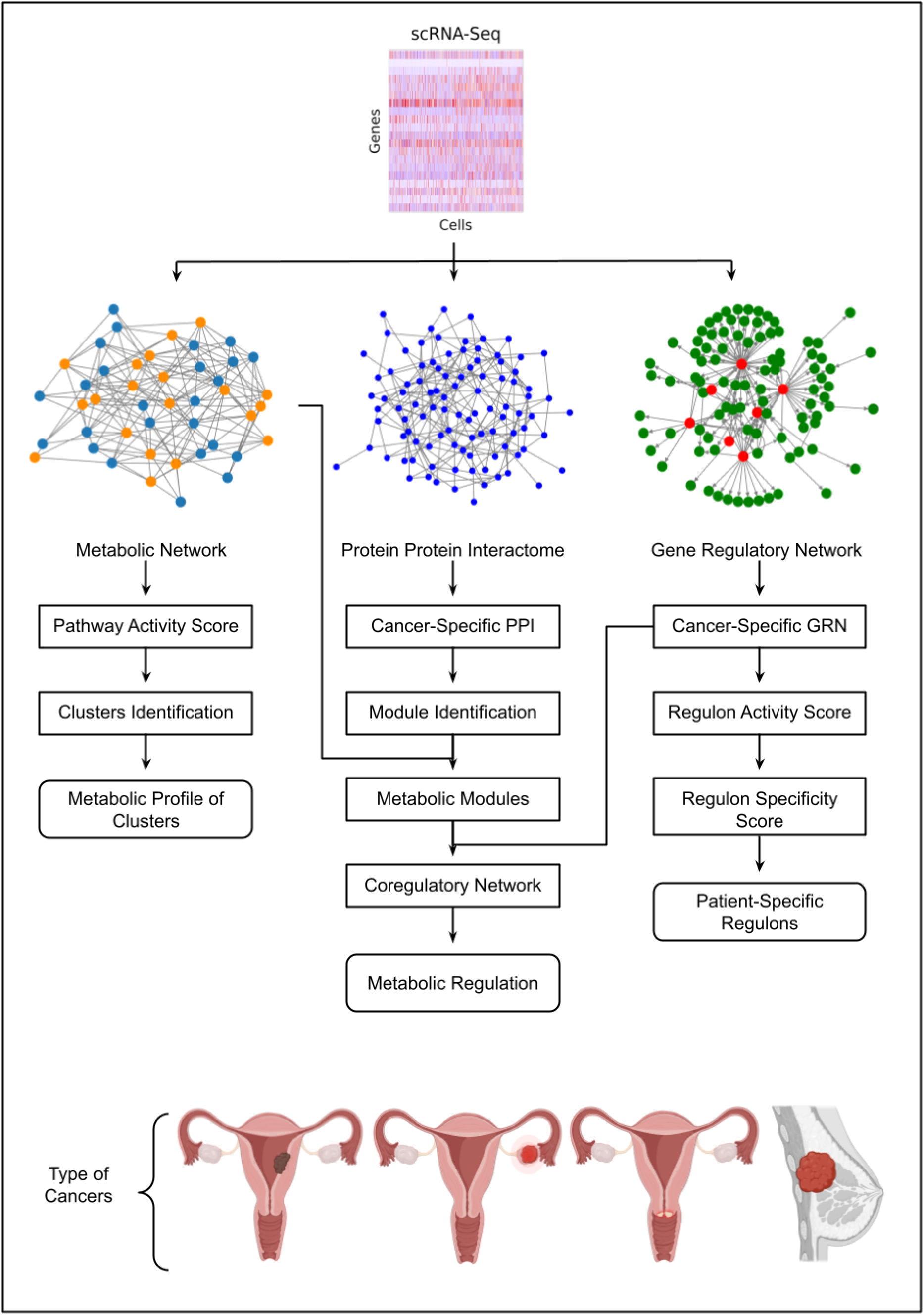
The pipeline for uncovering the metabolic regulation of gynecologic and breast cancers using a single-cell network approach (see methods for details).

## Results

### Heterogeneity of Metabolic Pathway in Cancer Cells

The metabolic pathways from the Human-GEM model were scored based on scRNA-Seq data to characterize malignant heterogeneity in gynecologic and breast cancers. We identified significantly overdispersed pathways that formed the basis for clustering malignant cells into distinct clusters (Figure 2A). We obtained 5 clusters of endometrial cancer, ovarian cancer, cervical cancer, ER+ breast cancer, and 6 clusters of TNBC (Figure 2 and 3). The chi-square test reveals a significant correlation between these clusters and patient information.

**Figure 2:**
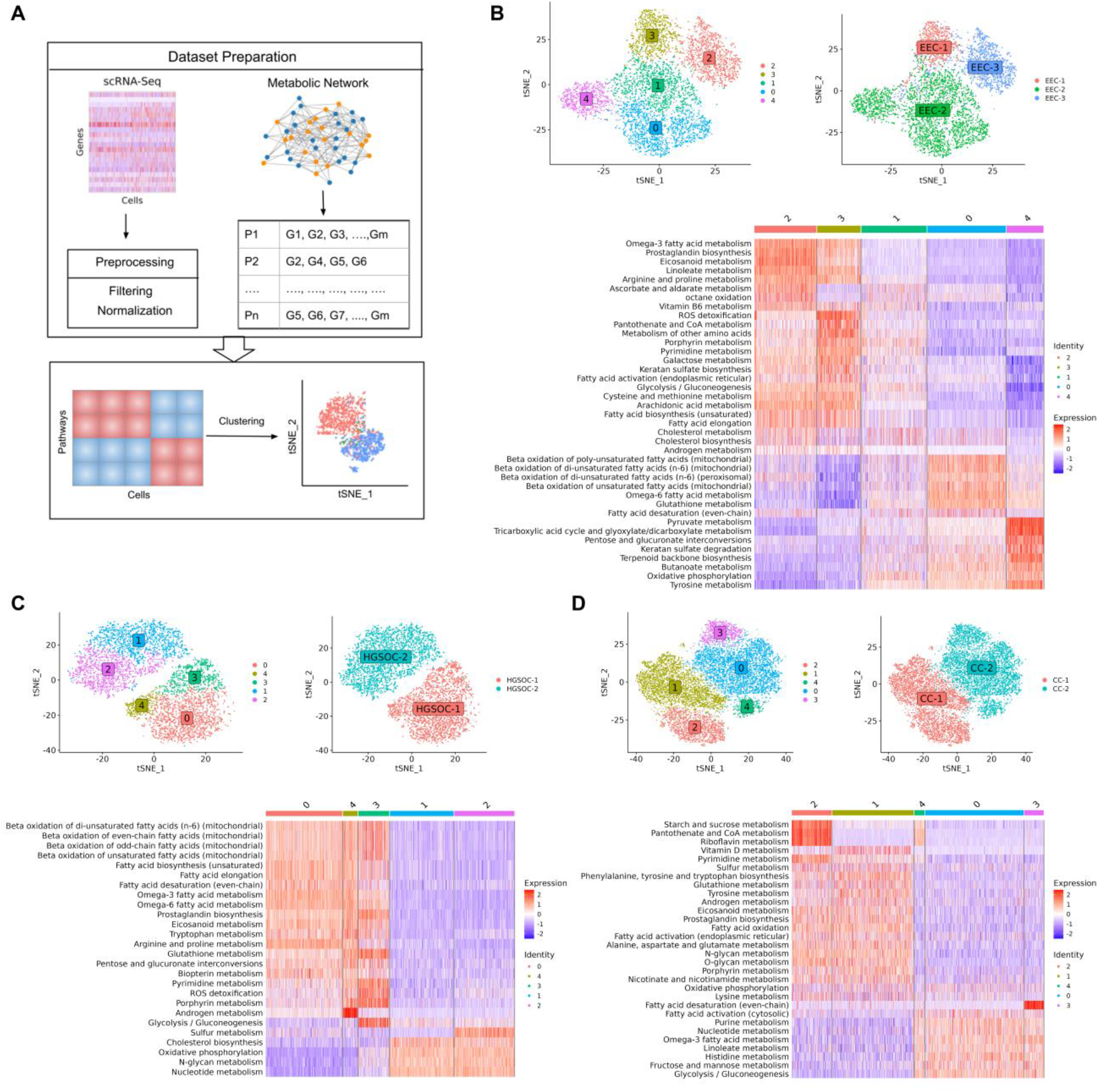
(A). Overview of the single-cell approach used to characterize metabolic heterogeneity, integrating the Human-GEM model with scRNA-seq data. The metabolic landscape of (B) endometrial cancer, (C) ovarian cancer and (D) cervical cancer, showing clustering of malignant cells, distribution of clusters across patients, and heatmap of metabolic pathway activity score.

**Figure 3:**
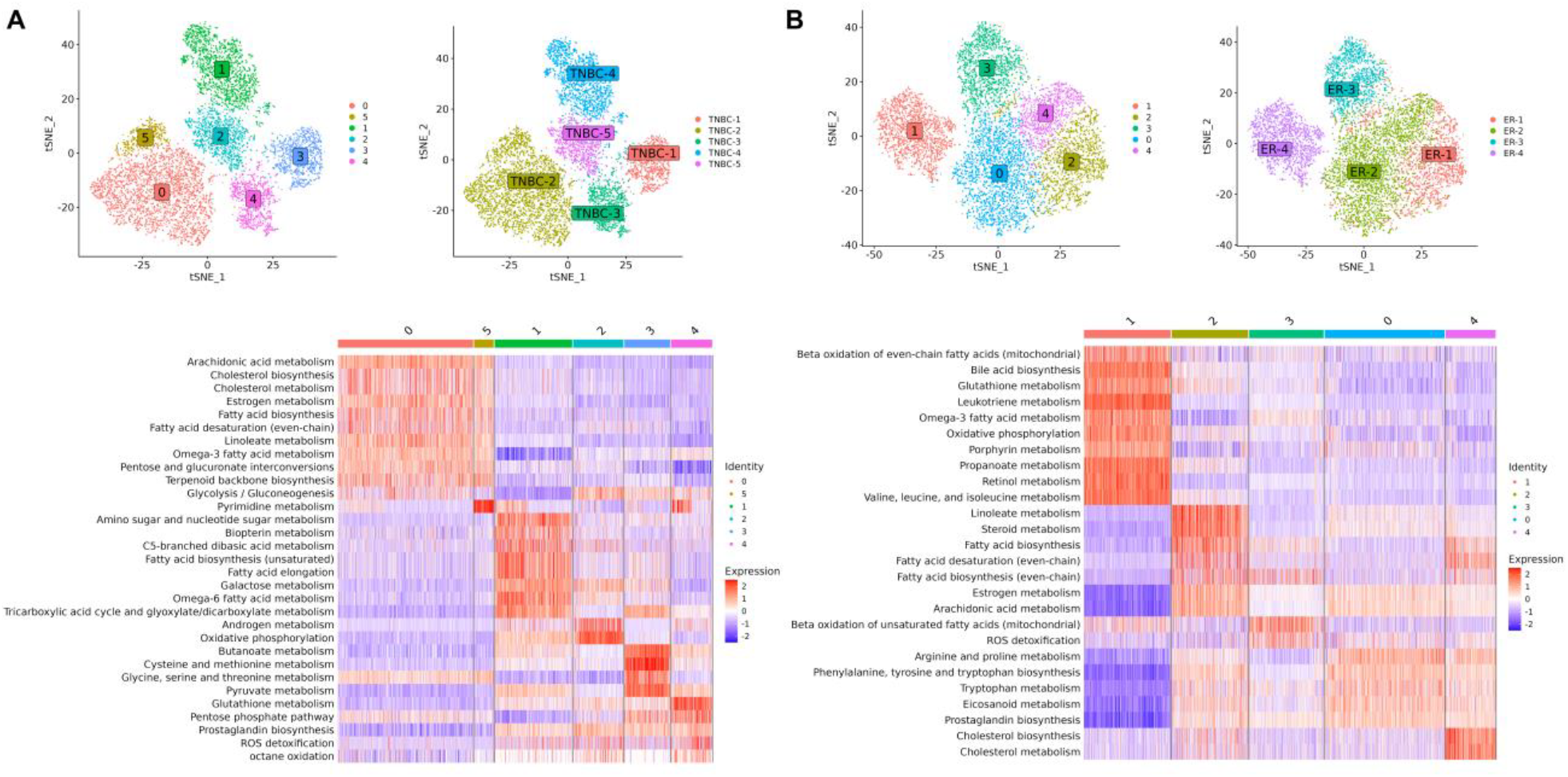
The metabolic landscape of (A) TNBC and (B) ER+ breast cancer, showing clustering of malignant cells, distribution of clusters across patients, and heatmap of metabolic pathway activity score.

In endometrial cancer, we observed a single cluster for patient EEC-1 (cluster 3) and EEC-3 (cluster 2), while EEC-2 displays three heterogeneous populations (cluster 0, 1, and 4) (Figure 2B). EEC-1 and EEC-3 share similar metabolic profiles with upregulation of glycolysis, fatty acid biosynthesis, androgen and cholesterol metabolism. However, differences in expression levels of some pathways were observed. EEC-1 exhibits elevated expression in pathways such as galactose metabolism, pantothenate and CoA metabolism, pyrimidine metabolism, and ROS detoxification. In contrast, EEC-3 shows high expression in pathways such as omega-3 fatty acid metabolism, prostaglandin biosynthesis, linoleate metabolism, and eicosanoid metabolism. In the case of EEC-2, we observed distinct metabolic profiles within its respective clusters. Cluster 0 displays high activity in fatty acid beta-oxidation pathways. Cluster 4 exhibits upregulation in energy metabolism pathways such as pyruvate metabolism, TCA cycle, and oxidative phosphorylation. Cluster 1 represents an intermediate state with mixed expression patterns across metabolic pathways.

In ovarian cancer, three heterogeneous clusters (cluster 0, 3, 4) emerged within HGSOC-1 and two clusters (cluster 1, 2) in HGSOC-2 (Figure 2C). Surprisingly, we observed a high expression of fatty acid metabolism, with simultaneous upregulation of both fatty acid synthesis and fatty acid oxidation in HGSOC-1, suggesting a possible complex interplay. Omega 3- and omega-6 metabolism is also highly expressed in HGSOC-1. In HGSOC-2, we observed oxidative phosphorylation, nucleotide metabolism, N-glycan metabolism, and cholesterol biosynthesis to be upregulated. Our findings align with recent studies indicating metabolic heterogeneity in ovarian cancer, particularly favoring oxidative phosphorylation in invasive migratory and cancer stem cells^11,12^. Cholesterol biosynthesis has been associated with enhancing the stemness properties of cancer cells, a crucial factor in initiating the metastatic cascade ^13^. Both ovarian cancer patients are characterized by intra-malignant heterogeneity. In HGSOC-1, cluster 0 shows high expression of arginine and proline metabolism. Cluster 3 has increased activity for pathways such as glycolysis, pyrimidine metabolism, glutathione metabolism, and ROS detoxification. Cluster 4 demonstrates upregulation in androgen metabolism.

In cervical cancer, we observed 2 heterogeneous groups (cluster 1, 2) in CC-1 and 3 groups (cluster 0, 3, 4) in CC-2 (Figure 2D). CC-1 is characterized by the upregulation of oxidative phosphorylation, androgen metabolism, amino acid metabolism, glycan metabolism, and eicosanoid metabolism. On the other hand, CC-2 displays high expression for pathways such as glycolysis, fructose and mannose metabolism, and nucleotide metabolism. In CC-1, cluster 1 shows upregulation in vitamin D metabolism, while cluster 2 has high activity for starch and sucrose metabolism, pantothenate and CoA metabolism, and riboflavin metabolism.

In TNBC, we identified one cluster per patient, with the exception of TNBC-2, which exhibits two clusters (cluster 0, 5) (Figure 3A). Cluster 1, 2, 3, and 4 corresponds to patient TNBC-4, TNBC-5, TNBC-1, and TNBC-3, respectively. TNBC-1 is characterized by the upregulation of one-carbon metabolism, including cysteine and methionine metabolism and glycine, serine, and threonine metabolism, crucial for nucleotide synthesis, methylation reactions, and antioxidant defense. In TNBC-2, both cluster 0 and cluster 5 share similar metabolic profiles. TNBC-2 shows upregulation in lipid metabolism, including cholesterol metabolism, estrogen metabolism, linoleate metabolism, arachidonic acid metabolism, omega-3 fatty acid metabolism, fatty acid biosynthesis, and glycolysis. TNBC-3 demonstrates elevated expression in pathways associated with redox homeostasis such as glutathione metabolism, pentose phosphate pathway, and ROS detoxification. The pentose phosphate pathway provides NADPH, necessary for converting oxidized glutathione (GSSG) back into its reduced form (GSH), which is vital for scavenging reactive oxygen species (ROS). TNBC-4 exhibits high activity in metabolic pathways, such as amino sugar and nucleotide sugar metabolism, galactose metabolism, and the TCA cycle. TNBC-5 displays upregulation of oxidative phosphorylation and androgen metabolism.

In ER+ breast cancer, we obtained three clusters corresponding to each patient except ER-2, which has two heterogeneous groups of malignant cells, cluster 0 and 4 (Figure 3B). Cluster 1, 2, and 3 corresponds to patient ER-4, ER-1, and ER-3, respectively. We observed the upregulation of lipid metabolism, including steroid metabolism, estrogen metabolism, and fatty acid biosynthesis in ER-1. The two clusters of ER-2 share a similar metabolic profile but exhibit differences in the expression levels of pathways such as arginine and proline metabolism, phenylalanine, tyrosine and tryptophan biosynthesis, which are upregulated in cluster 0, while cholesterol metabolism and fatty acid biosynthesis are upregulated in cluster 4. In ER-3, we observed that the estrogen metabolism is not expressed. ER-4 is characterized by the upregulation of oxidative phosphorylation, propanoate metabolism, glutathione metabolism, and valine, leucine, and isoleucine metabolism. However, estrogen metabolism is downregulated. Notably, ER-4 exhibits high metabolic activity compared to the other ER+ patients.

Overall, we found that malignant cell clusters of gynecologic and breast cancers show gene expression differences in cholesterol metabolism, eicosanoid metabolism, fatty acid metabolism, oxidative phosphorylation, glycolysis, amino acid metabolism, and redox pathways. The clusters also show differences in androgen and estrogen metabolism.

### Cancer Cell-Specific Protein-Protein Interaction Network

To uncover the connectivity pattern among genes in malignant cells, we aimed to infer malignant cell-specific PPI networks by integrating gene expression with PCNet. Network corresponding to gynecologic and breast cancers comprises approximately 14-25 % of PCNet nodes and 4-22% of PCNet edges (Figure 4A and Table S1). We observed that more than 50% of nodes and 30% of edges are shared between at least two cancers (Figure 4B & 4C). However, the cervical cancer network has more unique nodes and edges compared to the rest of the cancers. To better understand the network-level relationship among these cancers, we performed network proximity analysis. This analysis reveals that all gynecologic and breast cancers are significantly proximal to each other, with distance ranging from 0.4 to 0.8, except ovarian and cervical cancers (Z-score < -2 and p-value< 0.001) (Figure 4D). This suggests a shared relationship between gynecologic and breast cancers at the network level. Additionally, TNBC and ER+ breast cancer show the least proximity since both are subtypes of breast cancer. We also extracted a network common across gynecologic and breast cancers. This network consists of 140 genes and 580 interactions belonging to important pathways such as oxidative phosphorylation, estrogen response early, estrogen response late, and androgen response (Table S2).

**Figure 4:**
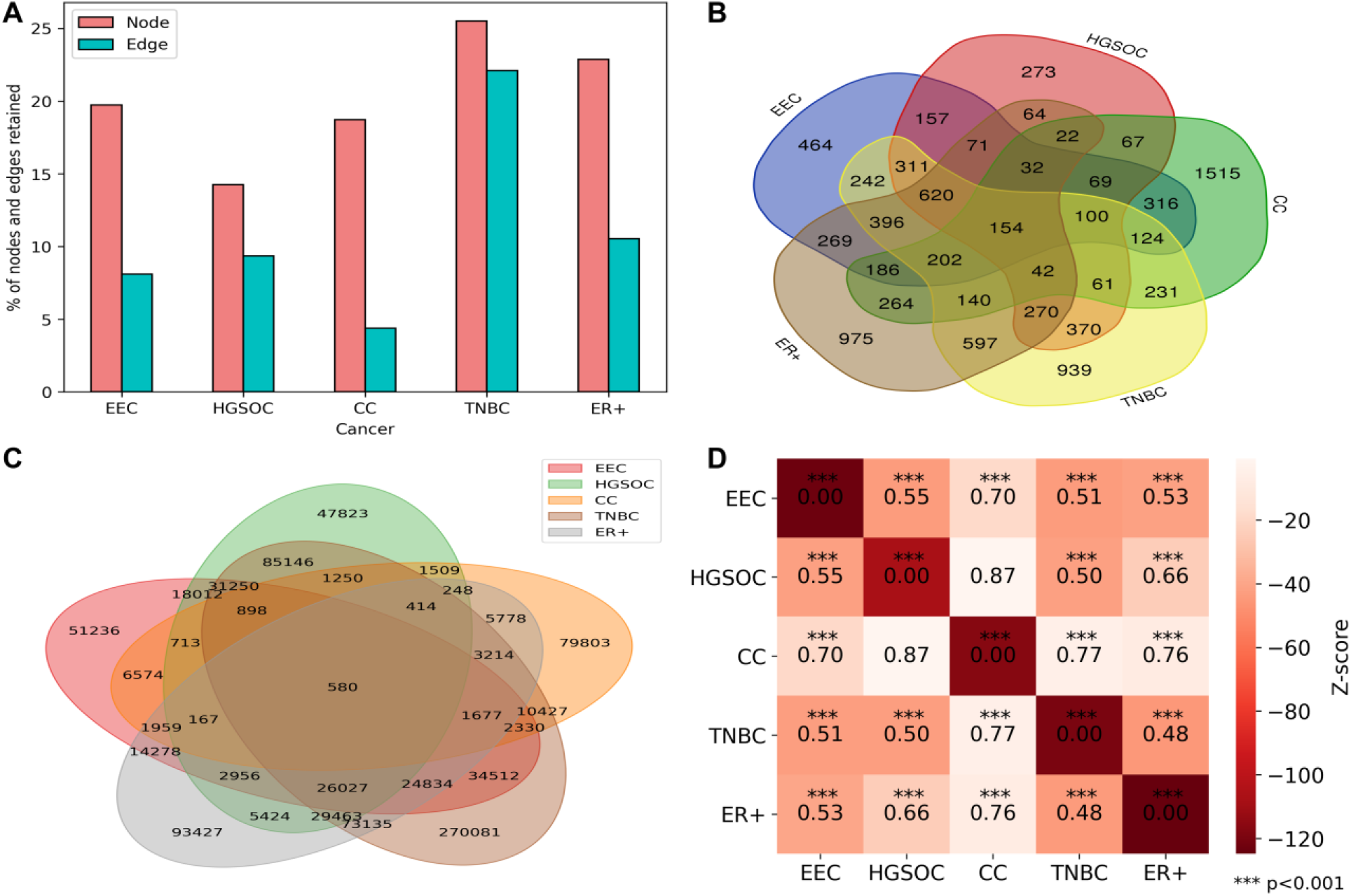
(A) Bar plot displaying the percentage of nodes and edges preserved in PPI networks of gynecologic and breast cancers. (B) Venn diagram illustrating the overlap of nodes in the PPI networks. (C) Venn diagram showing the overlap of edges in the PPI networks. (D) Network proximity analysis of gynecologic and breast cancers, with the heatmap indicating the distances between networks.

The PPI network is further used to identify metabolic modules, uncovering connectivity patterns among metabolic genes. Clustering was applied on each malignant specific PPI based on network topology. Subsequently, the pathway enrichment analysis of modules was performed to identify metabolic modules. We found two modules to be significantly enriched in metabolic pathways in endometrial cancer, ovarian cancer, and TNBC (Figure 5). For endometrial cancer, module M6 (EEC_M6) is enriched in lipid metabolism, while module M8 (EEC_M8) is enriched in oxidative phosphorylation, consistent with the pathway-level analysis. Similarly, for ovarian cancer, module M3 (OV_M3) shows enrichment in pathways such as cholesterol metabolism, omega-3 fatty acid metabolism, omega-6 fatty acid metabolism, arachidonic acid metabolism, and estrogen metabolism. On the other hand, module M5 (OV_M5) is enriched in oxidative phosphorylation. For TNBC, module M2 (TNBC_M2) is significantly enriched in beta-oxidation of fatty acids, TCA cycle, and oxidative phosphorylation. In contrast, the other metabolic module M10 (TNBC_M10) is enriched in cholesterol metabolism, glycine, serine and threonine metabolism, alanine, aspartate and glutamate metabolism, and phenylalanine, tyrosine and tryptophan metabolism.

**Figure 5:**
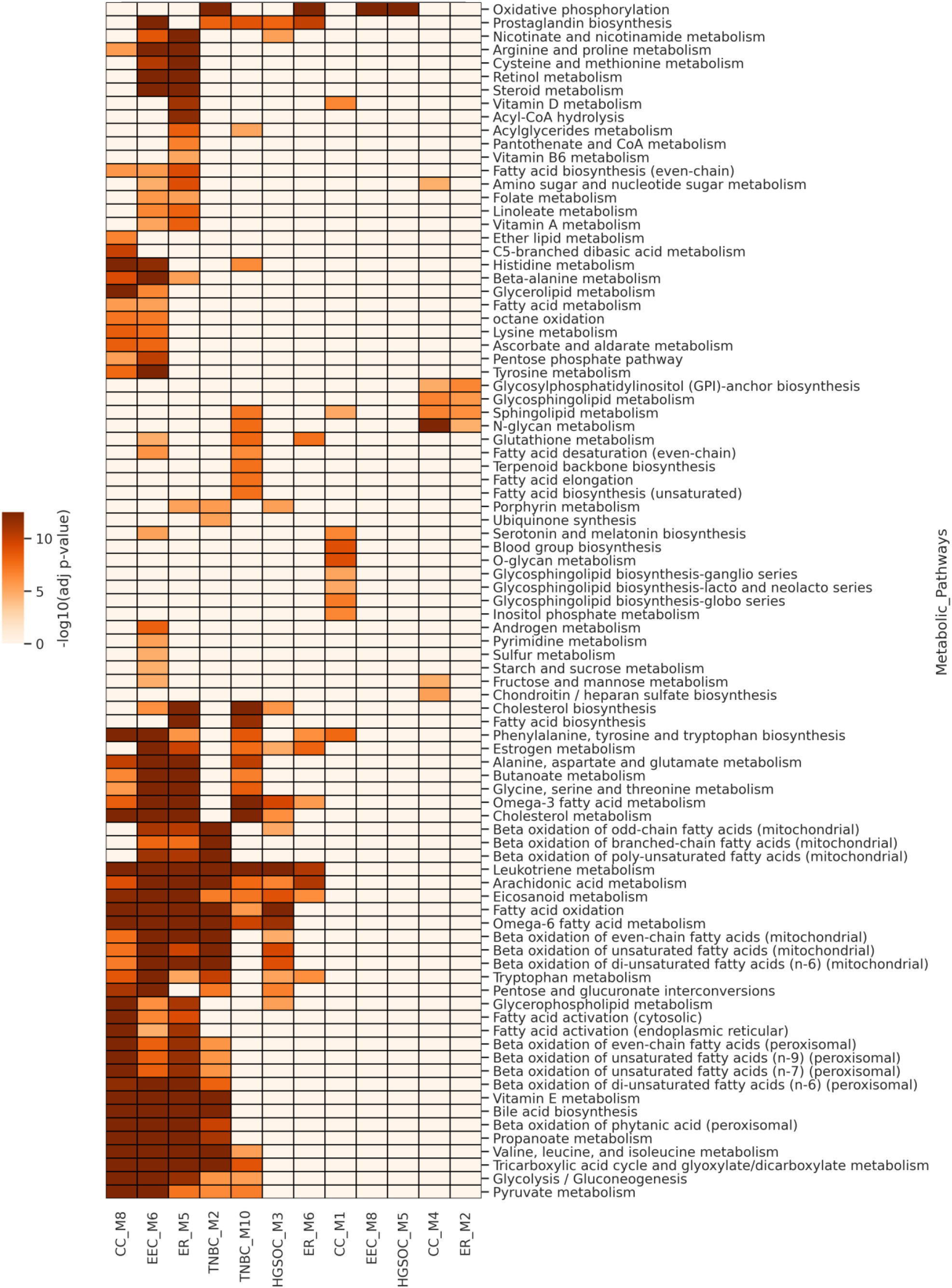
The heatmap showing the enrichment of modules associated with metabolic pathways identified from PPI networks of gynecologic and breast cancers.

In addition, we identified three metabolic modules for cervical and ER+ breast cancers (Figure 5). The M6 module in ER+ (ER_M6) is enriched in oxidative phosphorylation, similar to OV_M3, TNBC_M2, and EEC_M8 modules. The module ER_M5 also has similar overall enrichment profiles to the EEC_M6, TNBC_M2, and CC_M8 modules. Glycine, serine and threonine metabolism, and alanine, aspartate and glutamate metabolism are significantly enriched in EEC_M6 and ER_M5 modules. Other amino acid metabolism captured by this analysis include branched-chain amino acid metabolism and aromatic amino acid biosynthesis.

### Gene Regulatory Network of Gynecologic and Breast cancers

To investigate the transcriptional regulation mechanisms in malignant cells, we constructed gene regulatory networks and identified active regulons in gynecologic and breast cancers. The number of active regulons varied across different cancer types, ranging from 140 to 200 (Figure 6A and Table S3). Notably, we identified 31 common regulons across subtypes implicated in TNF-alpha signaling via NF-κB, hypoxia, G2-M checkpoint, and interferon-gamma response (Figure 6B and Table S4).

**Figure 6:**
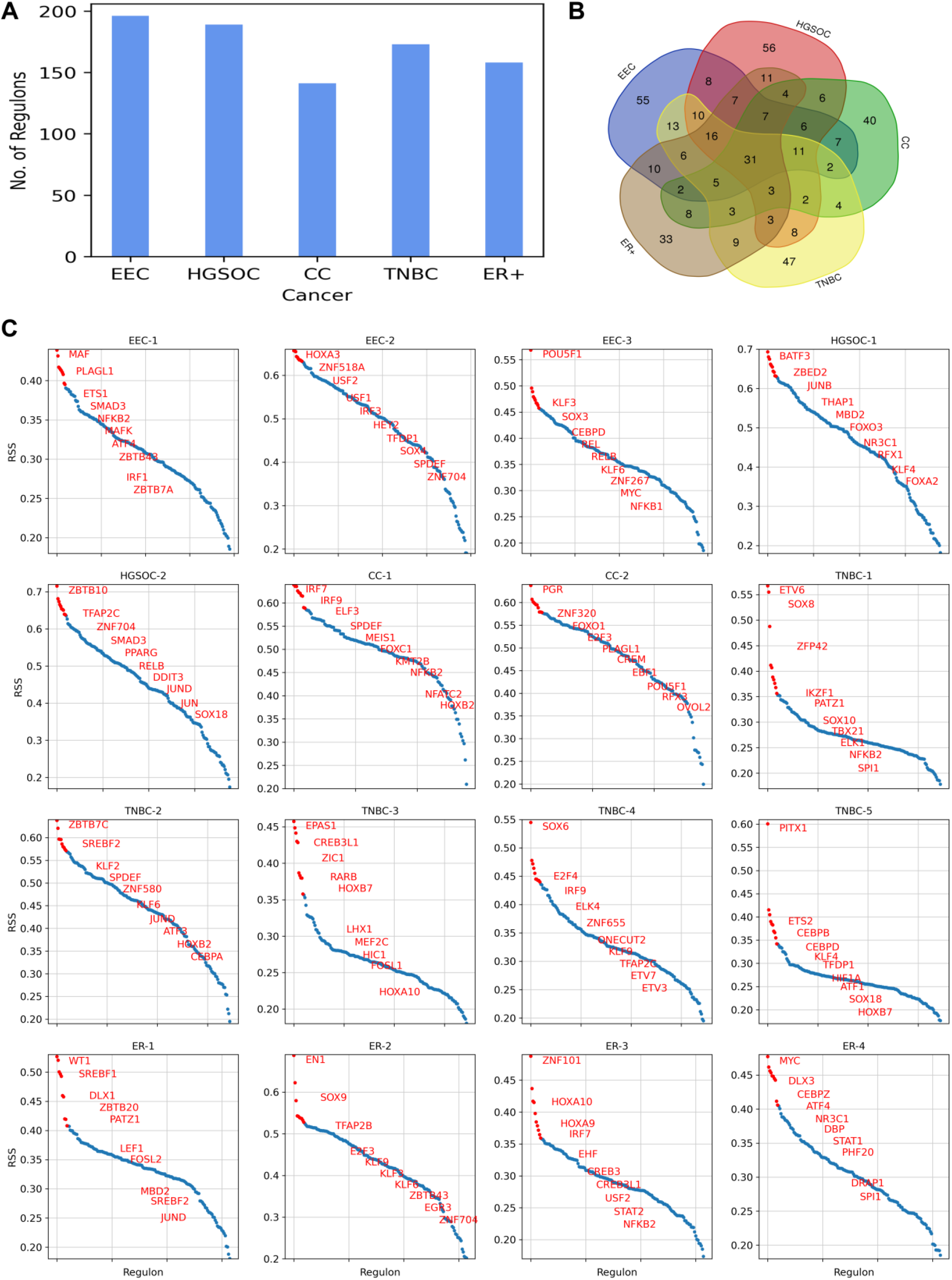
(A) Bar plot depicting the number of regulons identified in GRN of gynecologic and breast cancers. (B) Venn diagram illustrating the overlap of regulons among gynecologic and breast cancers. (C) RSS plot highlighting the top 10 patient-specific regulons in gynecologic and breast cancers.

To explore further, we characterized transcriptional regulation at the individual patient level and identified patient-specific regulons based on the regulon specificity score (RSS) (Figure 6C). In EEC-1, we found that the top regulons are involved in regulating the immune response and inflammation, such as regulons IRF1 and NFKB2 are critical for the activation of immune response genes, while ETS1 and SMAD3 contribute to the regulation of immune cell differentiation and function. Additionally, ATF4, a key regulator in the stress response pathway, is commonly expressed in EC cell lines, contributing to the tumor growth of endometrial cancer^14^. In EEC-2, the top regulons are involved in cellular differentiation, metabolism, and the cell cycle. The regulons HOXA3 and SOX4 play crucial roles in regulating gene expression during development, as well as in cell proliferation and differentiation. Additionally, the candidates USF1 and USF2 regulate the genes involved in lipid metabolism and glucose homeostasis^15^. EEC-3 is regulated by REL, RELB, and NFKB1, which are key players of the NF-κB signaling pathway, playing a vital role in immune responses, inflammation, and cell survival. KLF3 and KLF6, also associated with the immune system, are implicated in carcinogenesis^16^. Moreover, another candidate, POU5F1, is crucial for maintaining pluripotency in cancer stem-like cells and plays a pivotal role in tumor initiation and metastasis^17^.

In HGSOC-1, our analysis identified regulons that have been previously reported (Figure 6C). A recent pan-cancer analysis study highlighted the aberrant upregulation of ZBED2 in ovarian cancer and the enrichment of ZBED2-overexpressing cell lines in ovarian cancer-derived cell lines^18^. MBD2 expression is also significantly higher in high-grade serous ovarian cancer (HGSOC) tissue samples compared to normal tissue samples^19^. Another key regulon, FOXO3, is reported to be overexpressed in ovarian cancer cell lines^20^. The regulon NR3C1 encodes the Glucocorticoid receptor (GR), a hormone receptor involved in metabolic homeostasis and stress response. It accelerates glucose metabolism by driving stress response^21^. Similarly, in HGSOC-2, TFAP2C is reported to be upregulated in advanced ovarian carcinoma and is implicated in preventing ferroptosis, a regulated cell death process involving dysregulated iron metabolism, lipid peroxidation, and ROS accumulation^22^. Another important regulon, SMAD3, serves as a prognostic biomarker for ovarian cancer patients and plays a pivotal role in regulating the transforming growth factor-beta (TGF-β) pathway, which impacts metastatic processes in ovarian cancer^23,24^.

The regulons specific to CC-1 include IRF7, IRF9, NFKB2, and NFATC2, which are involved in the regulating genes associated with the immune response (Figure 6C). Another candidate, KMT2B, plays a role in regulating developmental genes and is found to be upregulated in cervical cancer cells and facilitates metastasis and angiogenesis in cervical cancer^25^. In CC-2, we identified PGR, which encodes a progesterone receptor. It is overexpressed in cervical adenocarcinomas, as reported by the TCGA study^26^. The regulon FOXO1, a key regulator of insulin signaling, controls metabolic homeostasis during oxidative stress. Other candidates, E2F3 regulates the cell cycle, EBF1 is crucial for B cell development and differentiation, and OVOL2 controls the epithelial-to-mesenchymal transition (EMT), which is critical for the development and cancer metastasis.

In TNBC-1, the regulons include ETV6, IKZF1, and SPI1, which are involved in the regulation of hematopoiesis and the development of blood cells (Figure 6C). Other candidates, SOX8 and SOX10, play crucial roles in regulating embryonic development and determining cell fate, while ZFP42 regulates pluripotency and differentiation in embryonic stem cells. In TNBC-2, we observed high expression of regulon SREBF2, which is involved in the regulation of lipid homeostasis, particularly cholesterol biosynthesis and metabolism. This observation aligns with our pathway-level analysis, which showed high activity in cholesterol metabolism for this patient. In TNBC-3, we observed high expression of the EPAS1 regulon, also known as hypoxia-inducible factor-2α (HIF-2α), which controls the transcription of genes involved in the cellular response to hypoxia and angiogenesis. Additionally, regulons CREB3L1 and HIC1 are involved in the cellular response to endoplasmic reticulum (ER) stress and DNA damage, respectively. In TNBC-4, high expression of TFAP2C was observed, consistent with the recent single-cell study of TNBC^27^. Other candidate regulons include E2F4, IRF9, ETV3, and ETV7. In TNBC-5, our analysis identified PITX1 as a highly expressed regulon. Overexpression of PITX1 has been reported in breast cancer cells, functioning as a critical regulator of estrogen receptor alpha-mediated transcriptional activity in breast cancer cells^28^.

In ER-1, the top regulon WT1 has been implicated in regulating EMT in breast cancer cells, with higher expression observed in the ER+ subtype^29^ (Figure 6C). High activity of candidate regulons SREBF1 and SREBF2 is associated with the upregulation of cholesterol metabolism in ER-1, similar to TNBC-2. Another key regulon, PATZ1, is involved in cellular proliferation and apoptosis and is overexpressed in breast tumors^30^. In ER-2, regulon TFAP2B is highly expressed and plays a crucial role in regulating tumor cell proliferation in breast cancer^31^. In ER-3, top regulons are linked to development (HOXA9, HOXA10), immune response (IRF7, STAT2, NFKB2), and stress response (CREB3, CREB3L1)). In ER-4, regulon MYC is highly expressed and regulates many cellular signaling and metabolic pathways. Its overexpression contributes to the development of resistance in ER+ breast cancer^32^.

### Crosstalk between Gene Regulatory Network and Protein-Protein Interaction Network in the control of cancer cell metabolism

To elucidate the underlying regulation of metabolic changes in gynecologic and breast cancers, we performed transcriptional factor enrichment analysis of metabolic modules identified from the cell-specific network. We found several transcription factors significantly enriched in these modules, highlighting their pivotal roles in metabolic reprogramming.

In endometrial cancer, we observed that XBP1, TFDP1, E2F1, RFX3, USF2, and FOXM1 target the metabolic genes in EEC_M6 and EEC_M8 modules (Figure 7A and S1A). XBP1 plays a critical role in the cellular stress response related to ER stress and regulates the transcription of genes involved in lipid metabolism, glucose metabolism, and immune responses^33^. It is upregulated in EC compared to normal tissues^34^. Also, high expression of E2F1 promotes proliferation, invasion, and migration of EC cells^35^. The candidate FOXM1 is overexpressed in endometrial cancer and promotes the proliferation, migration, and metabolic activity of EC cells^36^.

**Figure 7:**
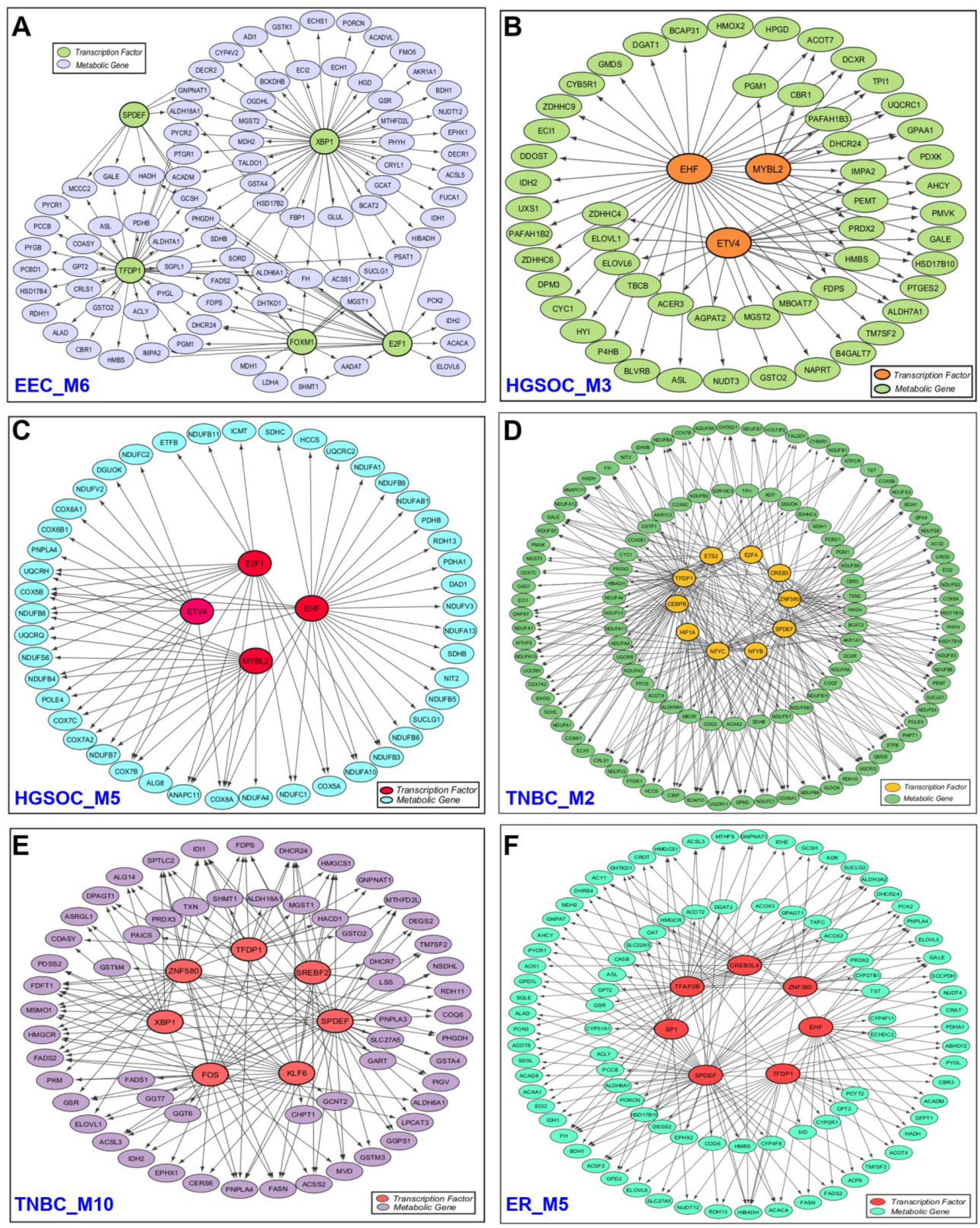
Interactions between significantly enriched transcription factors and genes within metabolic modules in gynecologic and breast cancers. For clarity, only the interactions between transcription factors and metabolic genes are displayed, while interactions between metabolic genes or between transcription factors are not shown.

In ovarian cancer, EHF, ETV4, E2F1, and MYBL2 target genes involved in lipid metabolism as well as oxidative phosphorylation (Figure 7B and 7C). EHF, a member of the E26 transformation-specific (ETS) transcription factor family, is overexpressed and associated with a shorter survival time in ovarian cancer^37^. It functions as a regulator of various metabolic processes and promotes cell proliferation^38^. Increased expression of ETV4 (ETS Variant Transcription Factor 4) contributes to chemoresistance in ovarian cancer^39^. A recent study also reported that MYBL2 promotes tumorigenesis in ovarian cancer^40^.

In cervical cancer, we identified one module (CC_M4) to be associated with transcription factors linked to ER stress response (CREB3L2, CREB3, and XBP1) (Figure S1B). CREB3 and CREB3L2, which are isoforms, also influence energy metabolism and cell differentiation^41^. Another candidate, CEBPD, is activated under hypoxic conditions and plays a role in promoting cancer cell metastasis and invasion^42^. The candidate HOXB2 is expressed in cervical tumor tissues and contributes to cancer progression^43^.

In TNBC, the important transcription factor is HIF1A, which targets genes of metabolic module M2 (TNBC_M2) (Figure 7D). HIF1A, a critical hypoxia-responsive transcription factor, is extensively regulated by inflammatory mediators and the hypoxic microenvironment in TNBC^44^. It targets multiple downstream genes to induce glycolysis, angiogenesis, invasion, metastasis, tumor stem cell enrichment, and immune escape, thereby promoting tumor proliferation and growth in TNBC^44^. Additionally, SREBF2 is associated with module M10 (TNBC_M10), targeting critical cholesterol and fatty acid metabolism genes, including HMGCR, HMGCS1, MVD, and FASN (Figure 7E). The candidates NFYB and NFYC are associated with module M2 (TNBC_M2). They are part of the trimeric protein complex NF-Y, which has been identified as a facilitator of SREBF activity and serves as a coactivator for sterol-regulated genes^45,46^. NF-Y also activates glycolytic genes and targets genes involved in various metabolic pathways^46^. The other candidates of TNBC are associated with ER stress response (XBP1, CREB3).

In ER+ breast cancer, all three metabolic modules showed an association with different transcription factors (Figure 7F, S1C, and S2D). The candidate SP1, enriched in module M5 (ER_M5), is involved in the estrogen signaling pathway. Estrogen receptors can interact with SP1 to regulate the transcription of target genes, which lack estrogen-responsive elements in their promoters^47^. This cooperative interaction helps in regulating the expression of multiple growth regulatory genes in breast cancer^48^. The candidate YBX1 is known to promote tumorigenesis, and its high expression is associated with poor prognosis in breast cancer patients^49^. Another candidate, MYBL2, enriched in module M2 (ER_M2), has been reported to drive cell cycle progression and influence EMT, significantly contributing to invasion and metastasis in breast cancer^50^. The other candidates are linked to cell differentiation (EHF, TFAP2B) and ER stress response (CREB3, CREB3L4).

Overall, the proposed approach captures the crosstalk between gene regulatory networks and protein-protein interaction networks that can drive metabolic reprogramming in gynecologic and breast cancers.

## Discussion

Metabolic reprogramming is a hallmark of cancer cells. Earlier scRNA-seq study showed significant variation in metabolic activity among different cell types within the tumor microenvironment^10^. In this work, we aimed to study the metabolic patterns and the regulation of metabolic pathways, which allow cancer cells to meet their increased energy and biosynthetic demands and adapt to the changing tumor microenvironment. Our study employed a comprehensive and systematic network-based approach to dissect the intricate landscape of metabolic regulation within malignant cells at the single-cell level. We first uncovered metabolic pathway profiles within malignant cells of gynecologic and breast cancers, elucidating key metabolic pathways and providing valuable insights into cancer metabolism. Through the inference of PPI and gene regulatory networks, we revealed the interactions among genes and the regulatory relationships between transcription factors and genes. Subsequently, we identified metabolic modules that highlight the connectivity patterns among metabolic genes. Finally, we pinpointed metabolic regulators by leveraging gene regulatory networks to identify transcription factors significantly enriched in metabolic modules. This analysis comprehensively captures the underlying regulation of metabolic changes in gynecologic and breast cancers, as the metabolic modules reflect gene connectivity patterns. This pipeline can also be applied to decipher the metabolic regulation of any disease at single-cell resolution.

We observed metabolic heterogeneity across different patents as well as within individual patient. In ovarian and cervical cancers, metabolic heterogeneity between patients can be correlated with their tumor stage differences. In contrast, metabolic variations are observed among EC patients even within the same tumor stage. The intra-patient heterogeneity in EC patient (EEC-2) indicates the presence of distinct tumor subpopulations with differing metabolic demands. Although all breast cancer patients in our study belong to either TNBC or ER+ subtypes, they also exhibited variation in metabolic profiles. This variability between patients suggests that various intrinsic (mutations, epigenetics) and extrinsic (tumor microenvironment) factors contribute to these differences^51^. TNBC patients can be subtyped based on the differences in glycolysis, oxidative phosphorylation, cholesterol metabolism, one-carbon metabolism, and redox metabolism. This extends the previous metabolic subtyping of TNBC into the lipogenic subtype, with upregulated lipid metabolism; the glycolytic subtype, with upregulated carbohydrate and nucleotide metabolism; and the mixed subtype, with major pathway dysregulation^52^.

Our study revealed that oxidative phosphorylation exhibits variation across gynecologic and breast cancers. This observation aligns with Xiao et al. (2019), who identified oxidative phosphorylation as a major contributor to metabolic heterogeneity in malignant cells^10^. Additionally, we observed variability in glycolysis activity and identified regulons associated with hypoxia, suggesting a probable interplay among oxidative phosphorylation, glycolysis, and hypoxic conditions within malignant cells. In endometrial and ovarian cancer, we observed mutually exclusive behavior with respect to oxidative phosphorylation and glycolysis. On the other hand, we observed a complex interplay of oxidative phosphorylation and glycolysis in TNBC, with one patient showing both processes to be upregulated (TNBC-5). This finding indicates that cancer cells exhibit metabolic plasticity, enabling them to switch between oxidative phosphorylation and glycolysis and occupy a hybrid state based on oxygen availability and metabolic demands. The hybrid glycolysis/oxidative phosphorylation state is associated with metastasis and therapy resistance^53^. Specifically, our analysis highlighted the elevated activity of hypoxia-inducible factors (HIF1A and EPAS1) in TNBC. The upregulation of HIFs in TNBC makes it resistant to different treatments^44^. We also found E2F1/4 and TFDP1 transcription factors to target oxidative phosphorylation in endometrial cancer, ovarian cancer, and TNBC.

We also observed variations in cholesterol and fatty acid metabolism and identified metabolic modules associated with these pathways at the network level. Cholesterol, primarily known for its role as a key component of cellular membranes, also serves as a precursor for the synthesis of steroid hormones and bile acids. Cholesterol metabolism involves de novo cholesterol biosynthesis, cholesterol uptake, efflux, and esterification^54^. Cholesterol biosynthesis is a complex, energy-consuming process starting with acetyl coenzyme A (acetyl-CoA) and proceeding through multiple subsequent reactions^54^. We observed alterations in cholesterol biosynthesis across gynecologic and breast cancers. Cholesterol can also enhance glycolysis in TNBC cells^55^. The regulation of cholesterol biosynthesis is primarily mediated by sterol regulatory element-binding proteins (SREBFs). Accordingly, SREBFs are expressed in breast cancer subtypes, TNBC and ER+. Interestingly, we observed that SREBF2 is directly targeted by the transcription factor KLF6 in TNBC, with both SREBF2 and KLF6 being enriched in the same metabolic module (TNBC_M10). This observation aligns with recent findings, suggesting that KLF6 may play a role in regulating SREBF2 and influencing metabolic gene expression in TNBC^56^. We also found that SREBF2 is expressed in endometrial and cervical cancers. Further, transcription factors EHF, ETV4, E2F1, and TFDP1 were found to control fatty acid metabolism in endometrial cancer, ovarian cancer, and TNBC.

We found SPDEF in our network-based analysis of TNBC and ER+ breast cancer (Figure 7 and S1). It is also highly expressed regulon in endometrial and cervical cancers (Figure 6). SPDEF is a member of the ETS (E26 transformation-specific) transcription factor family, which is expressed in hormone-regulated epithelia such as the prostate, breast, and ovary. SPDEF control genes related to fatty acid metabolism. It has been reported to be an oncogene in breast cancer and is upregulated during the progression of the disease^57^. Additionally, androgen and estrogen metabolism exhibit heterogeneous expression patterns within malignant cells of gynecologic and breast cancers. High levels of androgens have been reported to be linked with an increased risk of cervical cancer^58^. However, its role in endometrial cancer has been controversial^59^. Further, our study revealed a complex interplay between omega-3 and omega-6 metabolism in gynecologic and breast cancers. Omega-3 metabolism is important for the synthesis of anti-inflammatory eicosanoids (EPA and DHA), while omega-6 is involved in the synthesis of pro-inflammatory eicosanoids (arachidonic acid). We observed that in ovarian cancer, both omega-3 and omega-6 metabolisms are upregulated, while in endometrial cancer and TNBC patients, they are activated in a mutually exclusive manner. Arachidonic acid serves as the main precursor of several eicosanoid species, including prostaglandins and leukotrienes, which can modulate tumor microenvironment^60^.

Amino acid metabolism is altered in gynecologic and breast cancers, with distinct patterns observed across different cancer types. Arginine and proline metabolism exhibit heterogeneous expression patterns in endometrial, ovarian, and ER+ breast cancers. A recent metabolomic study of urine and serum samples from early-stage EC patients identified differential metabolites specifically involved in arginine and proline metabolism^61^. Another key pathway, one-carbon metabolism, is expressed in TNBC. This pathway is essential for supporting nucleotide and amino acid biosynthesis, epigenetic modifications, and antioxidant regeneration through the methionine cycle^62^. Additionally, variations in aromatic and branched-chain amino acid metabolism are observed in cervical and ER+ breast cancers. Furthermore, we observed differences in expression patterns of glutathione metabolism across gynecologic and breast cancers. Glutathione metabolism, ROS detoxification, and pentose phosphate pathway are highly expressed in some TNBC patients compared to others. This result indicates that cancer cells employ various strategies to manage oxidative stress, adapt to their environment, and enhance their survival. Through our integrated network analysis, we found that different transcription factors (XBP1, ATF4, CREB3L1, CREB3L2) associated with ER stress are involved in gynecologic and breast cancers. ER stress enhances the adaptability of tumor cells to unfavorable conditions and drives cancer progression. It affects glycolysis and lipid metabolism, which in turn promotes tumor growth, invasion, and metastatic potential^63^.

The gene regulatory network analysis uncovered significant activity of NF-κB-regulated regulons (NFKB1, NFKB2, REL, RELA, and RELB) in gynecologic and breast cancers. Nuclear factor-κB (NF-κB) is a family of stimuli-responsive transcription factors that control a wide range of genes involved in immune and inflammatory responses. This family includes five structurally related members, NFKB1, NFKB2, REL, RELA, and RELB, which regulate gene transcription by binding to a specific DNA sequence known as the κB enhancer, either as homo-or hetero-dimers^64^. NF-κB signaling pathway has been associated with cancer progression through its role in regulating EMT and metastasis^65^. Additionally, regulons involved in interferon-gamma response are expressed in gynecologic and breast cancers.

## Conclusion

The comprehensive analysis of scRNASeq data reveals the heterogeneity in metabolic processes among malignant cells in gynecologic and breast cancers. This heterogeneity highlights the complexity of metabolic reprogramming, which may play a crucial role in tumor survival, growth, and adaptation to the tumor microenvironment. Importantly, it uncovers the underlying regulatory mechanisms driving metabolic dysregulation, including the identification of key transcription factors and their metabolic targets, which together form an integrated network in these cancers. Our study provides a foundation for further investigation into the pathophysiology of cancer metabolism and developing targeted therapies that disrupt metabolic pathways critical to cancer progression and resistance.

## Methods

### Datasets

We obtained scRNA-seq datasets (feature-barcode matrices) for endometrial and ovarian cancer from the Gene Expression Omnibus (GEO) database under the accession number GSE173682^66^. The corresponding cell type annotations were retrieved from the provided supplementary files. The Broad Institute Single Cell Portal (https://singlecell.broadinstitute.org/single_cell) was used to download the scRNA-seq data with cell type annotations for breast cancer (Accession ID: SCP1039) and cervical cancer (Accession ID: SCP1950)^67,68^. The endometrial cancer dataset consists of five patients, all classified as having the endometrioid subtype with stage 1. The ovarian cancer cohort includes two patients, one at stage 2 and the other at stage 3. For cervical cancer, the dataset encompasses 7 patients, with 4 classifieds at stage 1 and 3 patients at stage 2. The breast cancer cohort comprises 26 patients: 11 ER+, 5 HER2+, and 10 TNBC.

To study cancer metabolism, we employed the latest version of the human genome-scale metabolic model known as Human-GEM (version 1.14.0)^69^. This comprehensive model encompasses 13,085 reactions, 8,499 metabolites, and 2,897 genes, providing a detailed representation of the standard metabolic processes within human cells.

### Data Preprocessing

The feature-barcode matrix for each patient was transformed into a Seurat object using the Seurat R package (version 4.4.0)^70^. We utilized the pre-filtered cell barcodes available in the supplementary/metadata files from the original dataset paper for downstream analysis. For endometrial and ovarian cancer datasets, quality control (QC) and doublet cell removal were conducted for each patient^66^. The QC metrics included three criteria: log-transformed UMI counts (> 2 MADs, low end), log-transformed number of expressed genes (> 2 MADs, low end), and log-transformed percentage of mitochondrial read counts (> 2 MADs, high end). Doublet cells were identified using both DoubletDecon and DoubletFinder. For cervical cancer datasets, the following QC criteria were applied for cell filtering: UMI counts < 8,000 or mitochondrial gene percentage < 10%, and gene counts between 500 and 4,000^68^. In breast cancer datasets, cells were retained if they exhibited a gene count exceeding 200, UMI counts surpassing 250, and a mitochondrial percentage below 20%^67^.

To ensure a balanced analysis without bias, we excluded the patients with a lower count of malignant cells (<1000) in endometrial, cervical, and breast cancer datasets. Consequently, for the downstream analysis, we retained 3 patients from endometrial cancer, 2 patients from cervical cancer, and 9 patients (4 ER+ and 5 TNBC) from breast cancer. Detailed information about patients is provided in the supplementary file (Table S5).

Next, we performed normalization using SCTransform, a variance-stabilizing transformation method that preserves true biological variation while removing the technical variability^71^. A prior study highlighted the better performance of SCTransform in comparison to alternative normalization methods for scRNA-seq analysis^72^.

### Calculation of Activity Score of Metabolic Pathways

To characterize the variation in the metabolic profile of cancer cells, we calculated the pathway activity score for each cell using Pagoda (pathway and gene set overdispersion analysis), a computational framework designed for characterizing transcriptional heterogeneity in scRNA-seq data. First, Pagoda fits a cell-specific error model using a modified mixture model approach. This approach models each gene in a cell as a mixture of amplification processes (negative binomial distribution) and dropout events (Poisson distribution). Next, it adjusts the variance of each gene by renormalizing the observed expression based on the error model estimates, which also includes batch corrections. To determine the significance of residual variance, the chi-square test is applied. Then, the method performs weighted Principal Component Analysis (PCA) for each gene set and utilizes the first principal component to quantify the activity of the gene set. Weighted PCA helps account for technical noise contributions. To assess the statistical significance of PAS, it compares the variance explained by the first principal component of the gene set with the expected variance calculated using random gene sets. It also corrects for multiple hypothesis testing to identify significant pathways with adjusted p-values < 0.05, termed as overdispersed, for subsequent analysis. We implemented this method using the Pagoda2 R package (version 1.0.11)^73^.

### Identification of Clusters based on Metabolic Pathways

To uncover the various metabolic states within cancer cells, we performed clustering on the pathway activity score data of metabolic pathways. First, we scaled the data to standardize the values, enabling a robust principal component analysis (PCA). We then computed a PCA matrix using the RunPCA function and selected appropriate principal components for subsequent clustering and visualization. We constructed a shared nearest neighbor graph with the FindNeighbors function, specifying 10 components, and applied the Louvain algorithm via the FindClusters function to identify distinct clusters. Finally, we visualized the clustering results using t-SNE plots, facilitating the interpretation of the metabolic landscape in cancer cells.

### Construction of Cancer-Cell Specific Protein-Protein Interaction Network

To gain a deeper understanding of the complex interactions among genes within cancer cells, we aimed to construct a comprehensive cancer-cell-specific PPI network. This approach allows us to decode crucial interactions and regulatory elements unique to cancer cells. We employed the SCINET method to generate cancer-specific PPI networks using scRNA-seq data of malignant cells^74^. SCINET integrates a global, context-independent reference interactome (parsimonious composite network (PCNet)^75^ with scRNA-seq data. The method involves robustly imputing, transforming, and normalizing scRNA-seq data to handle its inherent noise and sparsity. Then, it infers gene interaction probabilities at the cell level and defines group-level interaction strengths to build cell-type-specific interactomes. Subsequently, we applied the Leiden clustering algorithm to identify distinct modules within the network. To uncover the regulatory mechanisms governing metabolic processes, we performed hypergeometric tests to identify modules enriched in metabolic pathways and to determine the transcription factors associated with each module. An adjusted p-value threshold of < 0.01 was applied as the significance criterion for identifying metabolic pathways and transcription factors.

### Network Proximity Analysis

To quantify the similarity between different gynecologic and breast cancer subtypes, we performed a network proximity analysis using a network proximity metric defined as^76^:

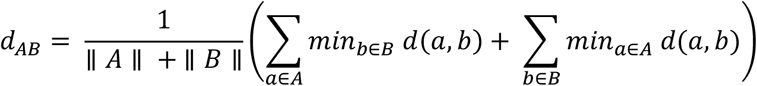

Here, *d*(*a, b*) is the shortest path distance between gene a from condition A and gene b from condition B within the interactome. To assess the significance of this distance metric, a Z-score was computed using a permutation test. This involved randomly selecting nodes from the entire network with degree distributions comparable to those of the nodes in the two sets. The Z-score was determined by performing 1,000 permutations as follows:

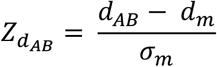

Here, *d*_*m*_ and *σ*_*m*_ are the mean and standard deviation of the permutation test. We conducted proximity analysis on gynecologic and breast cancers using their PPI network, with PCNet serving as the reference network. Two cancers were considered overlapping or significantly proximal in the interactome if they met the criteria of having a Z-score < *−* 2 and p-value < 0.001, indicating a degree of interaction similarity.

### Inference of Gene Regulatory Network

To elucidate the regulatory mechanisms underlying cancer progression at the single-cell level, we employed the SCENIC method to construct a gene regulatory network. SCENIC is a computational approach designed for inferring gene regulatory network from scRNA-seq data^77^. The pipeline consists of three main steps: first, it identifies potential regulatory modules by analyzing co-expression patterns among genes through a regression-based network inference method. Subsequently, it refines these co-expression modules by leveraging transcription factor motif information to remove indirect targets. Finally, the activity of these identified regulons is evaluated in individual cells using AUCell. Based on AUCell scores, we computed the regulon specificity score (RSS) to identify patient-specific regulons. This method was implemented using the pySCENIC (version 0.12.1) package available in Python.

